# Coupled neuro-fluid dynamics in human bioelectromagnetism

**DOI:** 10.64898/2026.10.08.757789

**Authors:** Tzvetan Popov, Gregory A. Miller

## Abstract

Electroencephalography (EEG) and magnetoencephalography (MEG) are widely interpreted as direct, non-invasive measures of neural population activity, assuming surrounding tissue and cerebrospinal fluid (CSF) act as passive volume conductors that distorts but do not contribute to signal generation. Using conductive-fluid phantom experiments, structural magnetic resonance imaging, and in vivo EEG and MEG recordings, the present report challenges this assumption. Mechanically driven conductive fluid generated structured, oscillatory, source-localizable electromagnetic signals resembling M/EEG in the absence of neural tissue, and interindividual differences in CSF volume systematically moderated M/EEG signal amplitude. These findings suggest that, at the scales accessed by invasive and non-invasive electrophysiology, electromagnetic and mechanical processes form a coupled system in which contributions of neural currents and fluid motion cannot readily be uniquely separated.

## Introduction

All waves propagate through a medium. In non-invasive electrophysiology, the physical environment through which brain waves travel is typically treated as a passive conduit rather than an active determinant of the measured signal. Electroencephalography (EEG) and magnetoencephalography (MEG) are commonly interpreted as direct manifestations of neural population activity, with cerebrospinal fluid (CSF), meninges, skull, and scalp treated and sometimes modeled primarily as sources of distortion rather than signal. A growing body of evidence complicates this view. Prominent low-frequency components of EEG and MEG covary strongly with physiological and mechanical processes such as head motion, respiration, gait, and cardiac dynamics, commonly treated as artifacts necessitating correction. Yet these “artifacts” can also be viewed as electromechanical phenomena in the medium through which brain waves and evoked waveforms propagate. Considering whether the medium itself contributes to electromagnetic recordings raises the question of the extent to which moving CSF may contribute to signal generation or modulation.

A particularly revealing challenge to standard EEG interpretation comes from posture-dependent effects. Rice and colleagues demonstrated that visually induced gamma-band responses systematically vary with participant posture despite identical sensory stimulation and task demands(1). Because the relevant perceptual computations remain unchanged, this finding implies that the electrophysiological correlate of a fixed psychological phenomenon is not invariant under changes in the physical configuration of the organism. Recent reports extend this concern. CSF-filled cavities exert a non-negligible influence on scalp EEG signals(2), non-neuronal sources can generate stable EEG patterns surviving standard preprocessing and source modeling(3), and combined EEG-fMRI studies demonstrate that large-amplitude CSF motion is phase-locked to cortical slow waves and sleep oscillations(4-6). Together, these observations raise the possibility that CSF contributions are not merely an unrelated or downstream consequence of neural activity but part of a coupled electromechanical process.

The present article contends that what unifies these findings is an underexamined property of the cellular environment itself: the presence of CSF as a distributed but continuous, highly conductive, mechanically dynamic compartment. CSF occupies sulci, fissures, cisterns, and perivascular spaces surrounding cortical and subcortical structures implicated in dominant electrophysiological rhythms. Critically, CSF is not static. It is continuously displaced by cardiac pulsation, respiration, posture maintenance, body movement, and even ocular activity, and it plays a central role in the glymphatic system(7, 8). Nevertheless, in EEG and MEG forward models CSF is typically treated as a passive, time-invariant conductor with a role limited to altering static field propagation.

From a physical standpoint, EEG and MEG do not measure individual neural events directly, but rather aggregate electric potentials and magnetic fields arising from closed neuronal current loops within the head(9, 10). Whereas neural transmembrane currents act as biological drivers of these loops, the pathways through which return currents close are determined by the geometry and mechanical state of the surrounding volume conductor. Changes in CSF distribution or motion therefore have the capacity to reshape macroscopic field patterns without altering microscopic neural firing. This issue is especially relevant for MEG, which is generally assumed to be less sensitive than EEG to volume-conductor effects and therefore more directly weighted toward tangential primary currents(11).

In summary, traditional EEG and MEG models assume that macroscopic signals arise solely from neuronal current dipoles closed by passive return currents in surrounding tissues(9, 10). This view, however, largely neglects the possibility that mechanically driven flows of cerebrospinal fluid - a moving ionic conductor - could themselves induce or modulate electromagnetic signals. By considering mechanically active CSF as part of the signal-generation loop, the present article investigates whether non-invasive electrophysiological signals reflect emergent properties of neural currents embedded in a mechanically active conductive medium.

Specifically, by combining phantom experiments, structural MRI, EEG, MEG, and mobile electrophysiology, we tested whether, as predicted by the foregoing arguments, CSF geometry systematically moderates electrophysiological signal amplitude and whether realistic, source-localizable electromagnetic patterns can emerge from conductive fluid dynamics in the absence of neural tissue. Across recording modalities and behavioral states, present findings indicate that CSF geometry systematically shapes measured signal amplitude and source structure, motivating a reframing of non-invasive electrophysiology as a coupled neuro-fluid measurement rather than a purely neural measurement.

## Methods

### Conductive-fluid phantom recordings

Electrophysiological recordings from the conductive-fluid phantoms were acquired using the mBrainTrain Smarting system (mBrainTrain LLC, Belgrade, Serbia). Signals were recorded using pre-wired sintered Ag/AgCl electrodes (1.5 mm diameter × 3 mm pellet; GreenTek, Wuhan, China). Data were acquired at 250 Hz. The same electrode type and amplifier configuration were used in all phantom recordings as in the human EEG experiments.

#### Beaker phantom

The cylindrical phantom consisted of a glass beaker (20 cm height, 14 cm inner diameter; total volume approximately 3.1 L) filled with conductive saline solution.

Electrodes were mounted in a custom 3D-printed scaffold approximating standard International 10-20 electrode locations and suspended within the phantom volume. Rhythmic fluid displacement was induced by manually squeezing and releasing a plastic pipette at 1 Hz rate positioned at predefined frontal or occipital locations relative to the electrode array, with the tip about 15 cm from the array.

#### Rubber-brain phantom

An anatomically realistic rubber brain model was submerged in conductive saline solution prepared as described above. Electrodes were positioned directly on the surface of the rubber brain at locations approximating standard International 10-20 electrode positions relative to the underlying anatomy. Electrodes remained fixed throughout the experiment while rhythmic fluid displacement was induced at 1 Hz using a stationary plastic pipette positioned at one of two predefined locations relative to the brain model. No biological tissue or active electrical source was present within the phantom. Source reconstruction of phantom recordings was performed using conventional EEG inverse procedures implemented in FieldTrip(12). Electrode positions were aligned to a standard three-shell boundary-element head model with stipulated conductivities of 0.33, 0.0041, and 0.33 S/m for brain, skull, and scalp compartments, respectively. Leadfields were computed using the boundary-element model, and source activity was estimated using linearly constrained minimum variance (LCMV) beamforming(13). The purpose of this analysis was to determine whether mechanically induced conductive-fluid signals could be assigned to plausible intracranial sources by standard EEG source-localization procedures rather than to provide a physically exact model of the phantom geometry.

#### Spherical biOrb phantom

The spherical phantom consisted of a biOrb® acrylic vessel with a volume of 105 L. The vessel was filled with water obtained from the host institution’s aquatic research facility, providing a naturally ionic aqueous medium routinely used for aquatic animal maintenance. Electrodes were attached to the inner surface of the spherical vessel.

Electrode locations were arranged approximately equidistantly along the vertical and horizontal meridians of the sphere, providing symmetric coverage of the conductive volume. The uppermost electrode (Cz) was positioned slightly below the upper opening of the vessel rather than at the geometric pole to ensure complete submersion within the conductive fluid and to avoid placement at the air–water interface. To induce fluid motion, the supporting table was displaced approximately 1 cm horizontally once by hand, producing free sloshing of the fluid within the sphere. Electrodes, cables, and recording equipment remained stationary during the subsequent oscillatory period. No biological tissue or active electrical source was present within the phantom. Source reconstruction was performed using a simplified single-sphere conductive-volume model implemented in FieldTrip. The model radius was set to 30 cm, with a conductivity of 1.5 S/m, corresponding to the conductive aqueous medium. A source model was constructed in FieldTrip by discretizing the conductive volume of the spherical phantom into a regular grid of dipole locations. Grid points falling outside the fluid-filled portion of the vessel were excluded, yielding a source space restricted to the conductive medium enclosed by the phantom. The source model was restricted to the fluid-filled portion of the sphere by truncating the model at the upper water surface and lower boundary of the vessel. Leadfields were computed for this truncated spherical source space, and source activity was estimated using LCMV beamforming. Power was estimated from virtual source time series in the dominant sloshing-frequency range and projected back into the spherical source space.

#### MEG fluid-displacement phantom

A 2 L plastic bottle filled with either saline solution (15 g sodium chloride dissolved in water; 0.128 mol/L) or deionized water was positioned inside the MEG dewar (whole-head MEG Triux, MEGIN Oy, Finland). Fluid motion was generated remotely through a 210 cm plastic tube connected to a 500 ml plastic bottle operated approximately 2 m from the sensor array. Rhythmic manual squeezing and releasing were done at 2 Hz while the operator listened to a metronome through the MEG intercom system. Identical pumping movements were performed in both saline and deionized-water conditions. No biological tissue or active electrical source was present within the phantom. Continuous MEG data were segmented into 30 s epochs and analyzed using multitaper frequency analysis with 0.15 Hz spectral smoothing in the 0.5-5 Hz frequency range. Planar gradient pairs were combined after spectral estimation. Saline-minus-deionized difference spectra and topographies were computed to isolate magnetic-field responses associated with fluid displacement.

### Human non-invasive recordings

#### Human EEG dataset

Human EEG data were reanalyzed from a previously published color-word Stroop experiment investigating the temporal dynamics of inhibitory control. Participants responded to the ink color of visually presented words while ignoring word meaning. The task included congruent, incongruent, and neutral trials and was designed to manipulate inhibitory-control demands. EEG was recorded using a 64-channel Ag/AgCl electrode montage and analyzed using source reconstruction and time–frequency methods. Full details regarding participant recruitment, experimental procedures, EEG acquisition, preprocessing, and behavioral analyses are provided elsewhere(14).

#### Human Connectome Project MEG datase

Resting-state magnetoencephalography (MEG) and structural MRI data were obtained from the Human Connectome Project dataset(15, 16). Continuous recordings were demeaned, detrended, segmented into 4 s epochs, and transformed into synthetic planar gradients(17). This transformation facilitates interpretation of sensor-level MEG data, because oscillatory power is maximal directly above the underlying source, yielding topographies that more closely reflect source location. No additional artifact rejection, ICA decomposition, or physiological signal correction was performed. T1-weighted images were acquired at 3 T using a high-resolution MPRAGE sequence with 0.7 mm isotropic voxels as part of the standard HCP imaging protocol. Detailed acquisition parameters are described elsewhere(16). Structural MRI segmentation, regional CSF quantification, participant stratification, power spectra analysis, and statistical analyses were performed as described above for the EEG dataset.

#### Welsh Advanced Neuroimaging Database MEG datase

MEG and structural MRI data were obtained from the Welsh Advanced Neuroimaging Database (WAND)(9). Whole-head MEG recordings were acquired at 1200 Hz using a 275-channel CTF axial gradiometer system. Structural MRI data consisted of high-resolution T1-weighted images acquired on a 3 T MRI system. Structural MRI segmentation, regional CSF quantification, participant stratification, and statistical analyses were performed as described above for the EEG dataset. Visual gamma-band responses were analyzed from the WAND visual task. In each trial, participants fixated centrally before viewing a circular sinusoidal grating that contracted toward fixation; after a variable interval, stimulus speed increased, and participants indicated detection of the speed change with a button press. MEG epochs were defined from -2 to 4 s relative to visual stimulus onset. Data were segmented into stimulus-locked epochs and demeaned. No additional artifact rejection, filtering, ICA decomposition, or physiological signal correction was performed. MEG data were transformed into synthetic planar gradients to facilitate interpretation of sensor-level power distributions. Time-frequency representations of power were computed using a Hanning-tapered sliding-window convolution approach. Frequencies from 2 to 100 Hz were estimated in 2 Hz steps. Window length varied as a function of frequency according to a constant five-cycle criterion (window length = 5/frequency), and spectral estimates were computed every 100 ms. Time-frequency analysis was applied to single-trial planar-gradient data. Auditory steady-state responses were analyzed from the auditory-motor task. MEG epochs were defined from -3 to 5 s relative to auditory stimulus onset. Data were segmented into stimulus-locked epochs, demeaned, and transformed into synthetic planar gradients. Time-frequency representations of power were computed using identical spectral parameters. In contrast to the visual gamma analysis, time-frequency decomposition was applied to the average over trials, thereby emphasizing phase-locked activity, and not to the single-trial data. In both visual and auditory tasks, statistical analyses were performed on the resulting raw power estimates without baseline normalization.

#### Mobile EEG navigation experiment

Human participants performed a delayed spontaneous alternation task adapted from the rodent literature(18). At the beginning of each trial, participants positioned their head in a chin rest and maintained fixation on a central fixation cross for 10 s while planning a forthcoming walking trajectory (see Supplementary Video). Following the delay period, a visual GO cue instructed participants to leave the chin rest, walk across the room, execute the planned turn, return to the starting position, and reposition themselves in the chin rest before initiating the next trial. The number of analyzable trials per participant was 86.4 ± 10.1 (mean ± SD; median = 87.5; range = 51-95). Variability in trial count primarily reflected intermittent loss of the synchronized accelerometer wireless data stream in a subset of participants or individual trials, resulting in exclusion of those trials from locomotor-frequency analyses. EEG was recorded continuously using a mobile 24-channel system (mBrainTrain Smarting) at a 250Hz sampling rate. Motion was recorded simultaneously using an Xsens DOT (https://www.xsens.com/wearables/xsens-dot) unit containing a triaxial accelerometer, gyroscope, and magnetometer sampled at 60 Hz. The sensor was attached centrally to the participant’s upper back at approximately chest level using an elastic strap. Accelerometer signals were streamed in real time via Lab Streaming Layer (LSL, https://labstreaminglayer.org/#/) and synchronized with EEG recordings. During the stationary delay period, binocular gaze position was additionally recorded using a Tobii Pro Fusion eye tracker but was not analyzed in the present study. The continuously recorded data were segmented into trial epochs extending from 0 to 26 s relative to trial onset. EEG data were rereferenced to the common average reference. Accelerometer recordings were resampled to the 250 Hz EEG sampling frequency and temporally aligned with the EEG data. For time-locked visualization of the data (Figure 3A), EEG signals were band-pass filtered between 0.5 and 5 Hz, and accelerometer signals were low-pass filtered at 5 Hz. Subject-specific averages were computed across all available trials. Cadence frequency was estimated from the vertical accelerometer component (AccelZ). Cadence power spectra were estimated using multitaper spectral analysis implemented in FieldTrip, employing discrete prolate spheroidal sequence (DPSS) tapers(19) with 0.05-Hz spectral smoothing. The dominant spectral peak of the accelerometer spectrum was identified for each participant and used as an estimate of individual cadence frequency. Participants were subsequently divided into slower- and faster-walking groups using a median split of the resulting cadence frequencies.

For source-level analyses, EEG data were projected into source space using LCMV beamforming similar to the procedures described in(14). A three-compartment boundary-element forward model (scalp, skull, and brain) based on the MNI152 template brain was used together with a standard electrode montage aligned to the template anatomy. Leadfields were computed for each location of a regular source grid covering the brain volume. Spatial filters were constructed from the electrode-level covariance matrix and applied to the EEG data to estimate source-level virtual-sensor time series at each grid location. Power spectra of virtual sensors were estimated as described above and parameterized using the specparam algorithm(20) to separate periodic and aperiodic spectral components. Source maps were generated from the resulting periodic spectral power estimates at the cadence frequencies identified from the accelerometer recordings. Source power estimates at individual grid locations were subsequently interpolated onto the standard MNI brain for visualization and group-level analyses.

### Data analysis

#### MRI segmentation and CSF quantification

Structural T1-weighted MRI scans were segmented using the SPM12 unified segmentation algorithm implemented in FieldTrip. Tissue probability maps for gray matter, white matter, and cerebrospinal fluid (CSF) were generated using the standard SPM12 tissue probability priors. Intracranial volume was estimated from the combined gray matter, white matter, and CSF probability maps. Total CSF volume was computed by summing CSF tissue probabilities within the intracranial compartment and converting voxelwise probabilities to milliliters. Regional CSF measures were obtained by projecting the Automated Anatomical Labeling (AAL) atlas(21) into individual MRI space and summing CSF probabilities within atlas-defined regions. Parietal CSF volume was estimated from AAL regions corresponding to the precuneus, postcentral, supramarginal, angular, and parietal cortices, whereas occipital CSF volume was estimated from occipital, cuneus, calcarine, and lingual regions. Subjects with visually apparent segmentation failures, characterized primarily by widespread misclassification of extracerebral fluid spaces as CSF, were excluded following quality-control inspection (N = 5). Volumetric analyses used the resulting parietal CSF and CSF fraction measures.

#### Spectral analysis

Oscillatory power spectra were computed from each artifact-free EEG epoch during the Stroop task, irrespective of trial condition, using multitaper spectral analysis implemented in FieldTrip. Spectral estimates were obtained using a Hanning taper and evaluated between 1 and 40 Hz. To separate oscillatory and aperiodic contributions, individual spectra were parameterized using the *specparam* algorithm(20). The periodic component was obtained by subtracting the fitted aperiodic spectrum from the original power spectrum. Participants were ranked according to parietal CSF volume, and those assigned to low (lowest 10%), intermediate (40–60%), and high (highest 10%) CSF groups were analyzed.

#### Time-frequency analysis

Time-frequency representations of power were computed separately for congruent and incongruent Stroop trials using a sliding-window convolution approach implemented in FieldTrip. Spectral power was estimated from 2 to 40 Hz using Hanning taper and 500 ms sliding window. Time-frequency estimates were computed from - 250 ms to 1.5 s relative to stimulus onset in 50-ms increments. Activity related to executive control was quantified as the difference between incongruent and congruent trials in each of low, intermediate, and high parietal CSF groups.

#### Statistical analyses

Inferential statistics used nonparametric cluster-based permutation testing(22) implemented in FieldTrip. To test whether electrophysiological activity varied monotonically with parietal CSF volume, participants were assigned ordinal predictor values according to their CSF group membership: low CSF = 1, intermediate CSF = 2, and high CSF = 3. At each channel, frequency, or time-frequency sample, FieldTrip’s independent-samples regression statistic was used to estimate the regression slope relating electrophysiological activity to CSF-group membership and to express this slope as a t-statistic. Cluster-level statistics were then computed from contiguous voxels exceeding the cluster-forming threshold, and statistical significance was assessed using Monte Carlo randomization following (22). This procedure tests for a monotonic linear trend across CSF groups while controlling the family-wise error rate across multiple electrodes, frequencies, and time points. For visualization and complementary model-based analyses, mean electrophysiological values were extracted from predefined sensor-frequency or sensor-time-frequency windows corresponding to the effects shown in Figures 2-3. Extracted values were entered into general linear models to quantify the association between electrophysiological amplitude and parietal CSF volume, while allowing inclusion of covariates such as gray-matter fraction, age, or sex where appropriate.

### Behavioral Stroop Performance

As previously reported(14, 23), behavioral performance showed a robust Stroop interference effect across participants, with longer reaction times and lower accuracy during incongruent (IC) than during congruent (C) and neutral (N) trials. To determine whether CSF volume was associated with behavioral conflict processing, reaction times during the Stroop interference for low, medium, and high CSF groups were compared. Although within each group incongruent trials differed significantly from congruent and neutral trials (all p < 0.001), the magnitude of these condition differences did not vary significantly by CSF group. Specifically, in one-way ANOVAs the three CSF groups were similar for the IC-C contrast (F(2,113) = 1.420, p = 0.246), the IC-N contrast (F(2,113) = 0.908, p = 0.406), and the C-N contrast (F(2,113) = 0.526, p = 0.593). However, the effects shown in Figure 2F reveal an important dissociation between electrophysiological signal strength and behavioral performance. Despite the three CSF groups’ similar behavioral Stroop, electrophysiological measures of Stroop interference, the mid-frontal theta, systematically scaled with CSF volume, reported below.

## Results

### Conductive Fluid Motion Generates Structured EEG Signals in the Absence of Neural Tissue

First, we examined whether macroscopic electrophysiological signals resembling neural EEG could arise in the absence of biological generators. A conductive fluid phantom consisting of saline in a cylindrical volume was instrumented with EEG-type Ag/AgCl electrodes arranged in standard EEG-like configurations (Figure 1A). Small, controlled mechanical perturbations were applied to the fluid via a plastic pipette pumped at a rate of 1 Hz, producing slow, rhythmic displacement without inducing electrode motion or cable artifacts. The resulting signals exhibited oscillatory structure with a dominant frequency at the pumping rate and were spatially specific to the site of mechanical perturbation (Figure 1A,B). To assess whether fluid-generated EEG-like signals can be distinguished from neural sources using conventional analysis pipelines, these phantom recordings were processed using standard preprocessing and source reconstruction procedures [e.g. (24)]. A boundary-element head model was employed(25), and inverse solutions were computed using beamforming procedures(13). Figure 1C,D illustrates that reconstructed sources localized consistently to focal regions within the modeled cortical space, producing spatial maps resembling those typically reported for neurophysiological oscillations or evoked responses. In a complementary conductive-fluid phantom procedure, a fluid-filled orb was placed on a table, and the table was gently displaced horizontally once to induce free sloshing of the fluid (Figure 1E). This single mechanical input produced sustained oscillatory activity at the natural sloshing frequency of the fluid. Importantly, the push itself had no spatially defined source within the orb volume. Nevertheless, the resulting oscillatory field showed a stable focal topography within the phantom (Figure 1F). This demonstrates that inverse solutions can assign mechanically induced fluid-current patterns to plausible internal sources even when the initiating mechanical event occurs outside the conductive volume. These results demonstrate that realistic cortical source estimates can emerge from conductive fluid dynamics alone.

**Figure 1:**
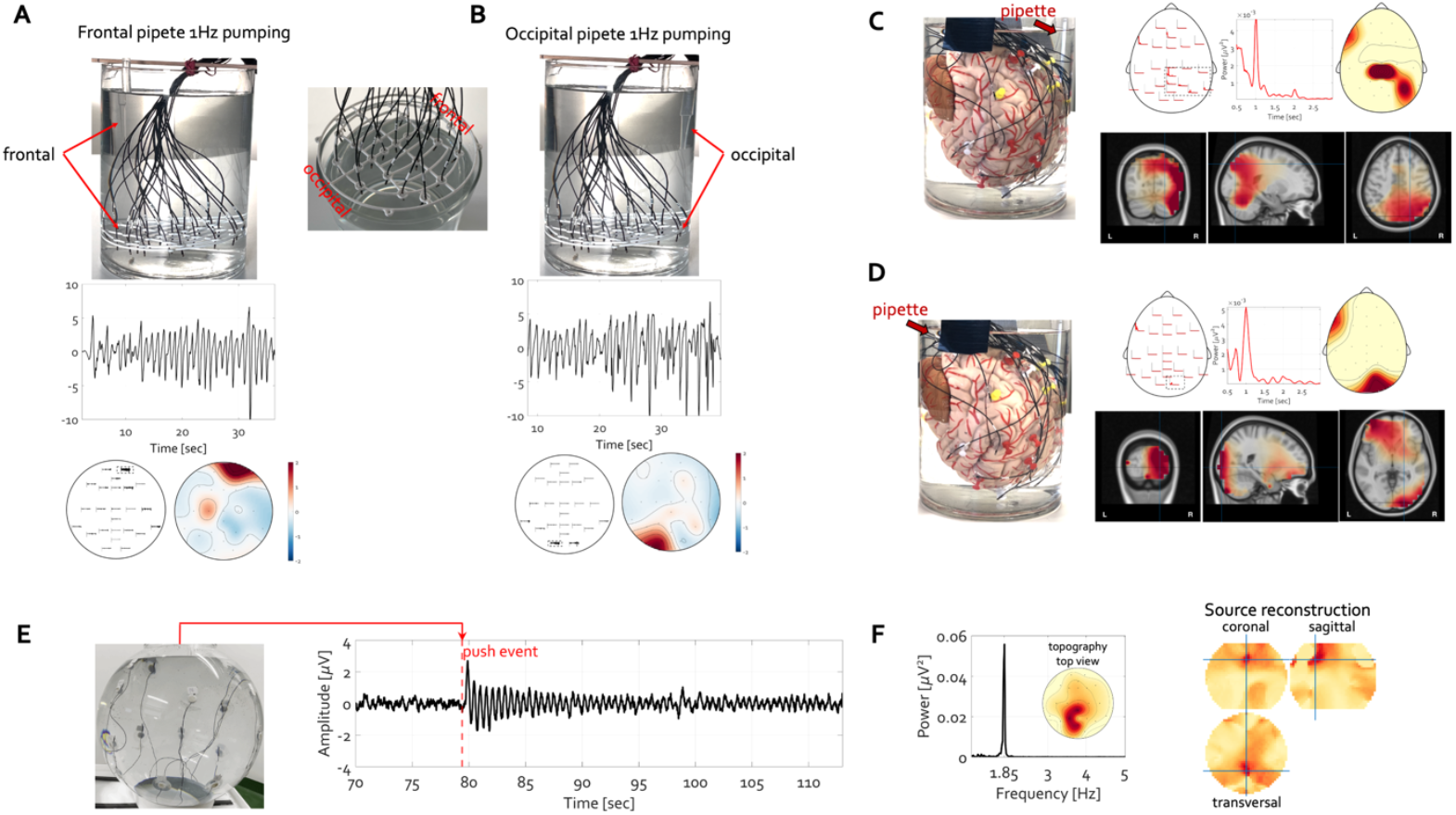
Mechanically driven conductive fluid generates structured, source-localizable electrophysiological signals in the absence of neural tissue. **A-B:** A conductive-fluid phantom was equipped with a scalp-like electrode array and filled with saline (saline prepared from tap water supplemented with approximately one teaspoon (5 g, 0.034 mol/L) of sodium chloride). Rhythmic fluid displacement was induced at 1 Hz by squeezing the 3 ml tip of a stationary plastic pipette positioned in either the frontal (A) or occipital (B) region of the phantom, relative to the electrode layout (top row, center inset). Despite the absence of neural sources, this very small water displacement produced clear oscillatory signals at individual electrodes at the bottom of the beaker (middle row; plotted electrode at lower arrow), with spatially distinct voltage topographies and reproducible field patterns (bottom panels) that depended on the location of mechanical input. **C-D:** An anatomically realistic rubber brain (frontal pole at bottom of beaker) was submerged in a similar conductive saline solution and instrumented with a scalp-like EEG electrode array. Periodic fluid displacement at 1 Hz was applied using a pipette positioned at different locations relative to the brain model (C: dorsal; D: ventral). Despite the absence of neural tissue, rhythmic vertical pipette motion produced clear oscillatory signals across electrodes (upper rows, left), with stable spectral peaks at the driving frequency (center) and distinct scalp voltage topographies (right). Using a standard boundary-element forward model and conventional inverse methods, these signals could be source-reconstructed to focal cortical regions corresponding to the location of the mechanical input (bottom rows; coronal, sagittal, and axial sagittal views overlaid on a template human MRI). **E:** A fluid-filled spherical phantom (biOrb®, 105 l) instrumented with scalp-like electrodes attached to its inner surface was gently displaced once by slightly pushing the supporting table, inducing free sloshing of the conductive fluid. The push event produced a transient deflection followed by sustained oscillatory activity in the microvolt range. **F:** Spectral analysis revealed a narrow peak at the dominant sloshing frequency (∼1.85 Hz) with a focal topography (top view). Despite the absence of a neural source and despite the external origin of the mechanical perturbation, conventional source reconstruction(13) using a single-shell spherical source model localized the resulting field to a focal region within the modeled volume.

### EEG Signal Amplitude Systematically Scales with Parietal CSF Volume in Young Adults

Second, we examined whether differences in CSF volume are associated with differences in EEG as resuls above would predict. Neurologically healthy young adults were stratified according to parietal CSF volume (Figure 2A) derived from individual high-resolution structural MRI. Across participants, EEG recordings acquired in the context of a color-word Stroop task revealed a robust relationship between CSF volume (Figure 2B) and electrophysiological signal amplitude (Figure 2D-F). The distribution of parietal CSF volume was split into deciles (low 0-10%, intermediate 40-60% and high 90-100%). Individuals with higher parietal CSF volume exhibited significantly higher alpha-band power (cluster-based permutation regression analysis, p < 0.05) and larger event-related brain potential amplitudes over posterior electrodes than those with smaller CSF volume. Importantly, this effect scaled monotonically across low, intermediate, and high CSF groups and could not be explained by differences in gray matter volume relative to total intracranial volume (Figure 2C). Several control analyses, including age distributions, Stroop, task accuracy, and condition-specific reaction-time contrasts across CSF groups, are provided in the Methods section and indicate that the observed electrophysiological scaling effects cannot readily be explained by CSF group differences in age or behavioral task performance.

**Figure 2:**
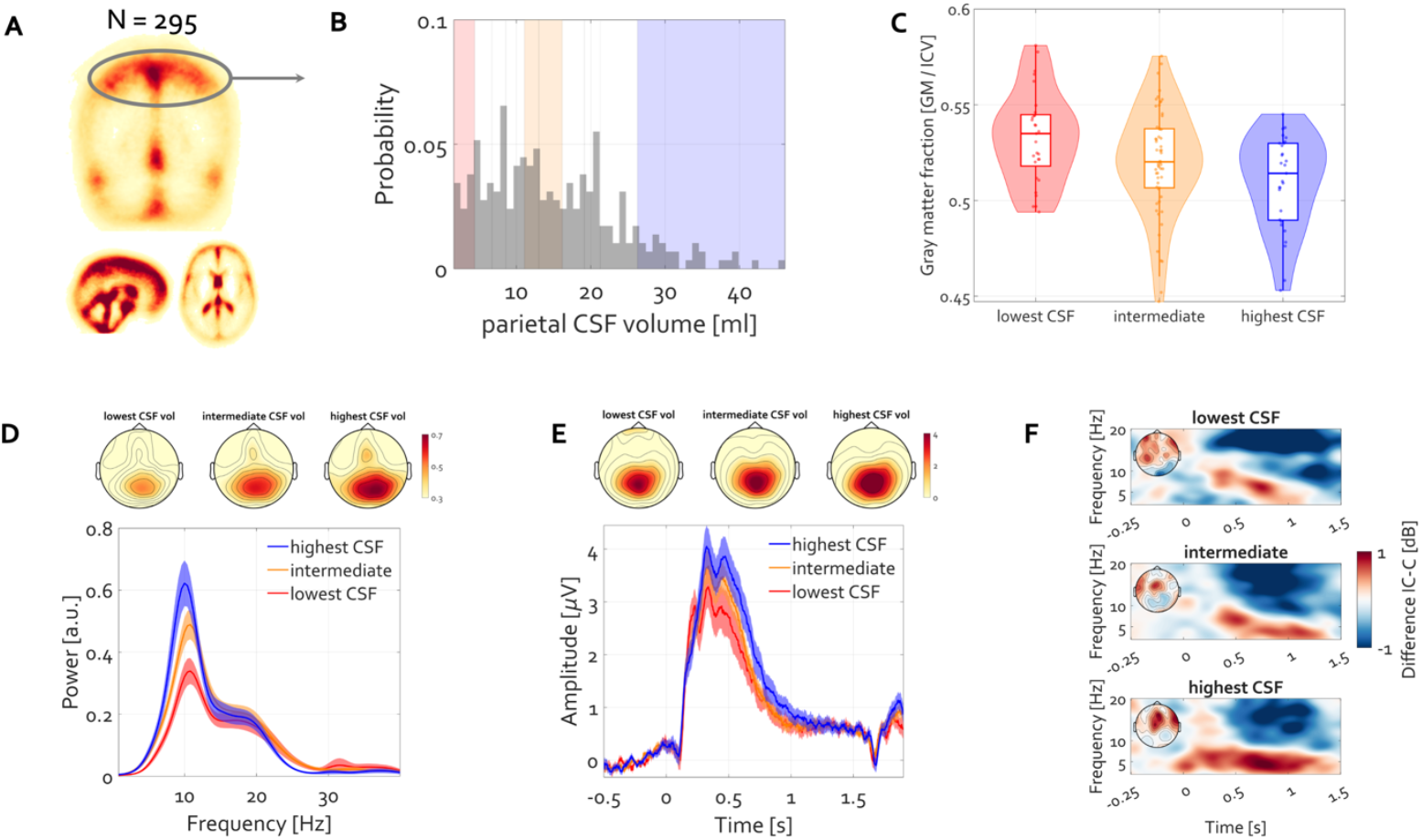
Cerebrospinal fluid volume systematically moderates EEG signal amplitude and spectral structure in healthy young adults. **A:** Group-level parietal CSF distribution derived from the structural MRIs of *N* = 257 healthy adults (150 female, ages 20-25; new analysis of dataset(14)). Parietal CSF volume denotes CSF probability volume within an AAL-defined parietal mask comprising Parietal, Precuneus, Postcentral, Supramarginal, and Angular parcels, resliced to individual anatomy and expressed in ml. Sagittal, axial, and coronal projections of CSF probability maps (% of whole-brain CSF per voxel), with the circled region indicating the parietal CSF compartment used for volumetric quantification. **B:** Histogram of individuals’ parietal CSF volume, with shaded regions indicating tertile-based grouping into low, intermediate, and high CSF volume cohorts. **C:** Distribution of gray matter fraction across participants grouped by parietal CSF volume. Gray matter fraction decreased systematically with increasing parietal CSF volume, such that participants in the highest CSF group exhibited the lowest gray matter fraction (ICV = intracranial volume). **D:** EEG signal properties stratified by parietal CSF volume (low, intermediate, high tertiles). Specparam-adjusted(20) power spectra showing systematic scaling of alpha-band power with CSF volume. Shaded regions indicate ±1 SEM across participants. 64-channel EEG data were acquired during an executive control (color-word Stroop) task(14) and collapsed across conditions for analyses shown here. **E:** Stimulus-locked event-related brain potentials (average reference montage), revealing monotonically higher peak amplitude with higher CSF volume despite identical task demands and similar behavioral performance. **F:** Time-frequency representation of power stratified by parietal CSF volume. Color scale indicates Stroop condition difference (incongruent IC minus congruent C) as change from baseline power in decibels. Topographical maps above or within each panel (D to G) depict corresponding scalp distributions. Across all measures, higher parietal CSF volume was associated with higher scalp signal amplitude and altered spectral characteristics.

Importantly, although behavioral Stroop effects remained stable across CSF groups, electrophysiological measures of Stroop interference systematically scaled with CSF volume. Increasing parietal CSF volume was associated with progressively stronger midfrontal theta activity during Stroop conflict processing (Figure 2F, cluster-based permutation regression analysis, p < 0.05).

### MEG Signal Amplitude Exhibits the Same CSF Dependence as EEG

These observations raise an important interpretational issue. EEG amplitudes are known to depend strongly on the conductive geometry of surrounding tissues, and the highly conductive CSF compartment may therefore amplify or spatially redistribute scalp potentials. Under standard EEG/MEG theory, such conductivity-related scaling should be markedly attenuated in MEG, which is generally assumed to be comparatively insensitive to volume-conductor effects(26-28). MEG therefore offers a critical test of whether the observed CSF dependence is limited to classical conductivity-related scaling of electric potentials or instead reflects a broader contribution of CSF dynamics to the macroscopic current loops underlying both electric and magnetic field measurements.

Third, we examined the sensitivity of MEG and EEG recordings to nonbiological “CSF” motion. To assess whether conductive-fluid dynamics alone can influence MEG measurements, a phantom was placed inside an MEG dewar (Figure 3A), a 2 l bottle filled with either saline solution (15 g sodium chloride, 0.128 mol/L) or deionized water. An experimenter approximately 2 m away from the dewar held a 500 ml plastic bottle connected to the 2 l bottle via a 210 cm plastic tube, listened to a metronome through the MEG intercom system, and pumped the 500 ml bottle at 2 Hz by repeatedly squeezing it. Because posture, muscle activity, and pumping movements were matched across saline and deionized conditions, movement-related contamination originating from the experimenter or recording environment would be expected to contribute similarly across conditions and to primarily affect sensors near the edge of the dewar.

**Figure 3:**
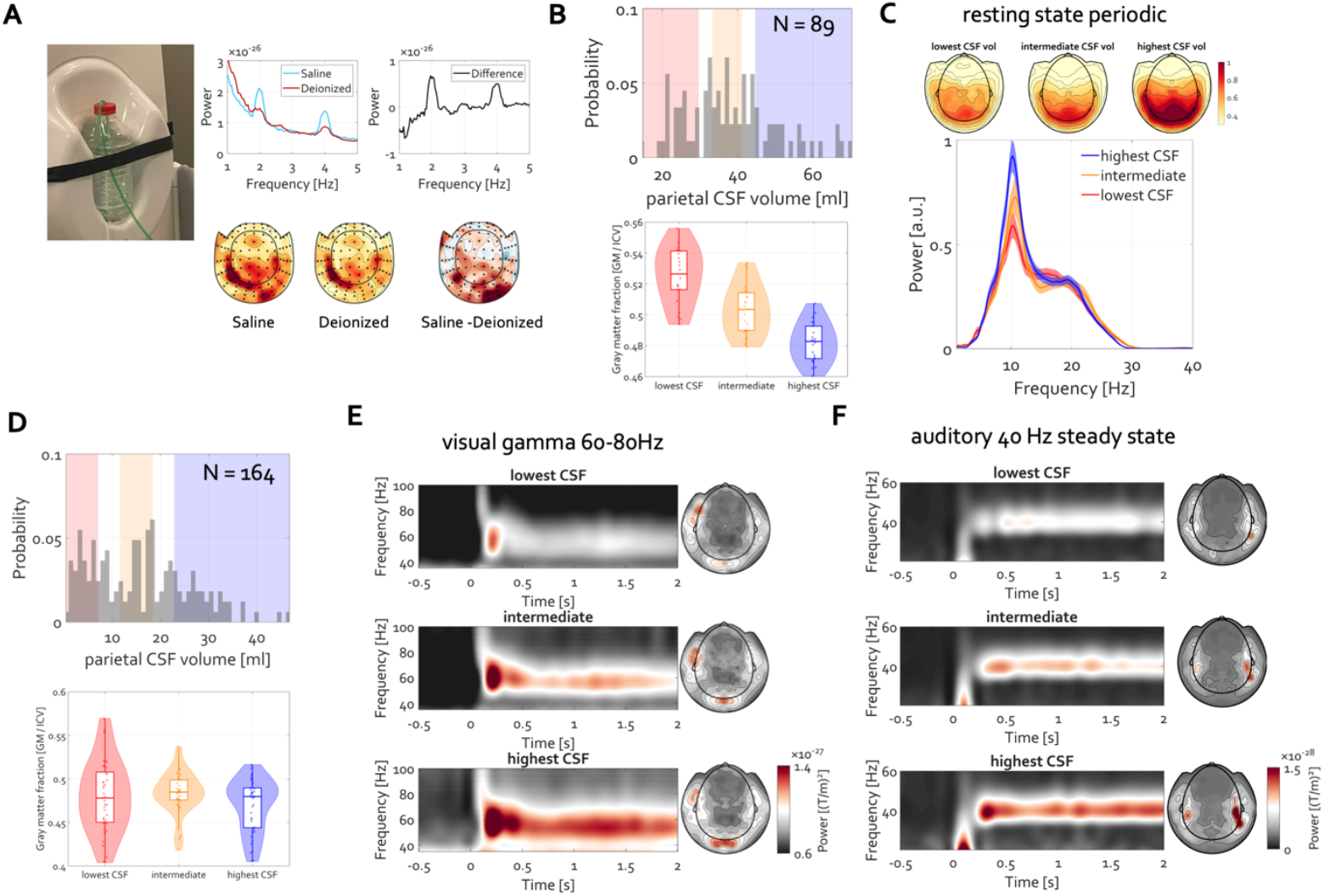
Cerebrospinal fluid volume systematically moderates MEG signal amplitude and spectral structure in healthy young adults. **A:** Conductive-fluid MEG phantom. Rhythmic input to saline produced stronger and more spatially structured magnetic field responses than deionized water, with spectral peaks centered at the pumping frequency and corresponding harmonics. Difference topographies revealed a left-parietal-dominant field distribution consistent with the position of the fluid-filled tube traversing the sensor array. **B:** *Top*- The histogram shows the distribution of parietal CSF volume across participants (N = 89) during resting MEG recording, with shading indicating stratification into low, intermediate, and high CSF volume groups. *Bottom*-Distribution of gray matter fraction across participants grouped by parietal CSF volume. Gray matter fraction decreased systematically with increasing parietal CSF volume, such that participants in the highest-CSF group exhibited the lowest gray matter fraction. **C:** MEG results stratified by parietal CSF volume (low, intermediate, high tertiles). Specparam-adjusted posterior sensor power spectra showing systematic scaling of alpha-band (8-14 Hz) power with CSF volume. Shaded regions indicate ± SEM across participants. Topographical maps depict corresponding scalp distributions. **D:** As in B but for the N = 164 cohort of the WAND dataset. **E:** Visual gamma-band responses (50-70 Hz) elicited by moving visual grating as a function of parietal CSF volume. Time-frequency representations of raw power (no baseline adjustments) are shown separately for participants with high, intermediate, and low CSF volumes. Corresponding sensor topographies are displayed to the right of each panel. Visual gamma-band activity tracked with parietal CSF volume, with stronger oscillatory responses in participants with larger CSF volume. **F:** Auditory steady-state responses elicited by 40 Hz auditory stimulation as a function of parietal CSF volume in the WAND dataset. Time-frequency representations of power (no baseline adjustment) are shown separately for high, intermediate, and low CSF groups, with corresponding sensor topographies displayed to the right. Auditory activity tracked parietal CSF volume, again with stronger oscillatory responses in participants with larger CSF volume.

Despite these matched conditions, saline pumping produced stronger and more spatially structured magnetic field responses than deionized water, with spectral peaks centered at the pumping frequency and corresponding harmonics (Figure 3A). Difference topographies revealed a left-parietal-dominant field distribution consistent with the position of the fluid-filled tube, which traversed the sensor array centrally with slight leftward parietal bias. These findings demonstrate that mechanically driven conductive-fluid dynamics alone can generate measurable, spatially organized magnetic field perturbations detectable with MEG sensors in the absence of neural tissue.

To further assess whether the relationship reported in sections above was specific to electric potential recordings or extended to magnetic field measurements, the same CSF-volume analysis was applied to publicly available MEG datasets with accompanying structural MRI(15, 29). Replicating the EEG findings, Human Connectome Project MEG data revealed a robust monotonic association between parietal CSF volume and signal amplitude during resting-state recordings (Figure 3B-C). Participants with larger parietal CSF volumes exhibited significantly higher posterior alpha-band power (8–12 Hz) and altered aperiodic activity (cluster-based permutation regression analyses, p < 0.05). Importantly, participants with the largest CSF volumes exhibited the strongest alpha-band signals despite having the smallest gray matter volumes, arguing against a simple source-to-sensor distance confound. Additional demographic and gray matter control analyses are provided in the Supplemental Material.

In a second MEG dataset(29), participants underwent visual gamma-band and auditory 40-Hz steady-state paradigms. Replicating the resting-state MEG findings, participants with larger parietal CSF volumes again exhibited systematically larger neuromagnetic responses, including stronger occipital gamma-band activity and bilateral temporal auditory steady-state responses (Figure 3D-F). Gray matter fraction differed only weakly across CSF groups (see Supplemental Material).

A final analysis tested whether the CSF-alpha relationship generalized across datasets and modalities, with regression analyses on a pooled sample of the EEG(14), HCP MEG(15), and WAND MEG(29) cohorts (total N = 546, 256 female). Parietal CSF volume, gray-matter fraction, age, sex, and dataset identity were entered simultaneously as predictors of alpha-band amplitude. The full model explained 27% of the variance in alpha amplitude, *F*(6, 539) = 33.40, *p* = 2.72 × 10^−34^. Parietal CSF volume was positively associated with alpha amplitude before partialling out the other predictors, *r* = 0.425, *p* = 2.51 × 10^−25^. The magnitude of this association initially differed across datasets, as indicated by a CSF × Dataset interaction, *F*(2, 540) = 3.80, *p* = 0.023. After partialling out gray-matter fraction, age, and sex, the CSF × Dataset interaction was no longer significant, *F*(2, 537) = 2.25, *p* = 0.107. Thus, although the magnitude of the CSF-alpha association showed some heterogeneity across datasets, this did not survive accounting for demographic and anatomical differences. Criticially, parietal CSF volume remained a positive predictor of alpha amplitude after partialling out variance associated with gray-matter fraction, age, sex, and dataset identity, Δ*R*^2^ = 0.031, *b* = 0.00622, SE = 0.00130, *t*(539) = 4.79, *p* = 2.18 × 10^−6^.. Results were qualitatively unchanged when alpha amplitudes were z-scored within each dataset before pooling. This convergence of EEG and MEG results across substantial samples supports the present contention that CSF geometry shapes the macroscopic current loops responsible for both electric and magnetic field measurements, consistent with a shared dependence on the physical structure of the conductive medium.

### Movement- and State-Dependent Low-Frequency Signals Cluster Around CSF-Rich Regions

Fourth, we examined the contribution of organism locomotion to the fluid-motion phenomena reported above. The EEG and MEG datasets examined thus far were acquired under typical stationary recording conditions. In natural behavior, by contrast, cerebrospinal fluid is continuously displaced by posture maintenance, locomotion, heart beats, respiration, and head acceleration. As neuroscience increasingly expands to include wearable and mobile electrophysiology in naturalistic settings, these mechanically induced CSF dynamics become particularly relevant. Locomotion represents perhaps the most extreme scenario for large-scale rhythmic CSF displacement, as each heel strike and phase of the gait cycle generates repetitive inertial and pressure-related perturbations throughout the cranial cavity. Mobile EEG recordings acquired during walking therefore provide a stringent test case for evaluating whether mechanically driven CSF dynamics can generate structured, source-localizable electrophysiological signals resembling canonical neural activity.

Anatomical and neuroimaging studies indicate that CSF-related parasagittal dural spaces are not uniformly distributed across the cranial cavity but instead exhibit a pronounced posterior-parietal predominance(30, 31), with recent ultra-slow flow MRI demonstrating comparatively elevated and highly reliable net CSF flow dynamics within parasagittal frontal, sensorimotor, and posterior parietal regions surrounding the superior sagittal sinus(32). We therefore predicted that gait-related electrophysiological components would localize preferentially to central-parietal regions, where CSF volume is greatest, and express dominant frequencies matching individual gait pace.

Participants performed a natural walking task while mobile EEG and accelerometry were recorded. Sensor-level data revealed clear rhythmic modulation of EEG signals time-locked to walking onset (Figure 4A), with oscillatory activity emerging at frequencies closely matching individual gait frequency (Figure 4B). Source reconstruction revealed that peak power localized predominantly to central sensorimotor and posterior parietal regions, with peak voxel locations differing systematically as a function of walking speed (Figure 4C,D). ANOVA revealed strong main effects of Group and Condition, as well as a robust Group × Condition interaction (all p < 0.0001; Figure 4F).

**Figure 4:**
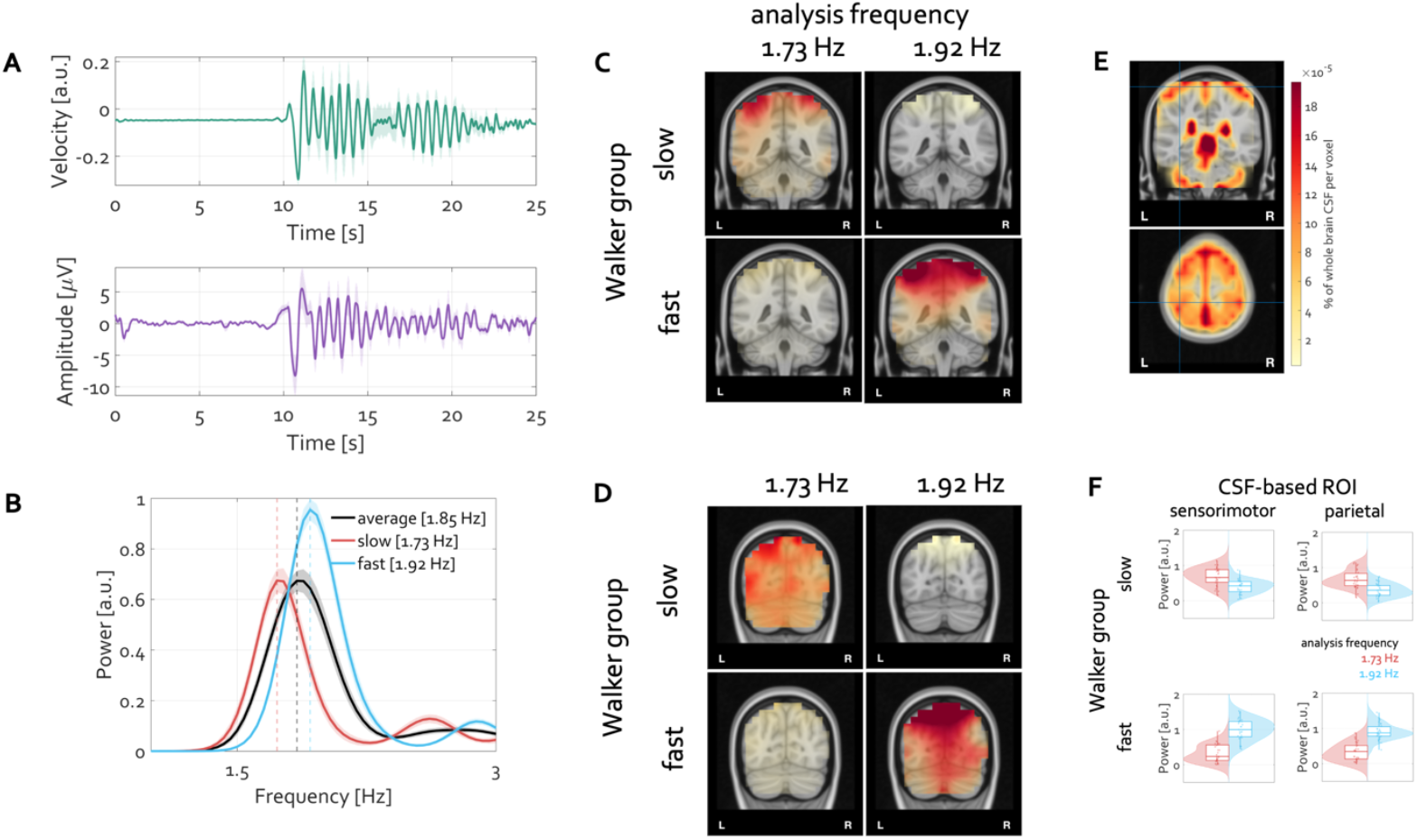
Gait-locked low-frequency EEG localizes to CSF-rich sensorimotor and parietal regions and scales with walking frequency. **A:** Group-averaged accelerometer velocity (top, arbitrary units) and occipital EEG signal (bottom, µV, average of O1 and O2) time-locked to gait onset, from N = 54 healthy adults age (22.9 ± 3.9, 38 female). Shaded regions indicate ± SEM across participants. Following a stationary 10 s wait, a prominent low-frequency oscillation emerged in both kinematic and EEG signals during locomotion. **B:** Power spectra of EEG source signals averaged across sensorimotor and parietal regions of interest (ROIs) centered at left parietal MNI (-11 -65 70), right parietal MNI (7 -65 70), left sensorimotor MNI (-38 -38 70), and right sensorimotor MNI (34 -38 70), showing a clear peak aligned with individual gait frequency. Participants were split into slow and fast walkers based on mean stepping frequency (slow: 1.73 Hz; fast: 1.92 Hz), with EEG peak frequency closely tracking walking speed. **C, D:** Coronal slices (y = -44) including central sensorimotor cortex from source reconstruction maps (via LCMV beamformer) for slow walkers (top rows) and fast walkers (bottom rows) showing EEG power at the two groups’ peak gait frequencies (left columns: 1.73 Hz; right columns: 1.92 Hz). For each group, maximal source power localized preferentially to central sensorimotor cortex. Notably, slow and fast walkers’ peak power emerged at their respective gait frequencies (upper left and lower right subpanels). **D: Same as C but for parietal cortex (y = -74). E:** Distribution of cerebrospinal fluid (CSF) probability across 257 participants from an independent structural MRI cohort, warped to a standard brain(14). CSF volume was maximal over dorsal parietal and midline regions in the supine MRI position, providing an anatomical reference for regions more susceptible to fluid-mediated effects. **F:** Quantification of gait-frequency-locked source power extracted from CSF-informed sensorimotor and parietal ROIs. Boxplots show that slow and fast walkers exhibit systematically different power distributions aligned with their respective gait frequencies, consistent with a movement-dependent mechanism rather than a fixed neural oscillation.

Although these signals are traditionally classified as motion artifact, source reconstruction localized them consistently to regions containing or adjacent to large CSF compartments. The frequency of these oscillations matched walking rhythm rather than canonical neural oscillatory bands, yet their spatial organization and reconstruction mirrored those of accepted neural signals. These findings extend the observed CSF dependence of EEG and MEG signals from static laboratory conditions to dynamic, naturalistic behavior characterized by continuous mechanical perturbation of cranial cavity.

## Discussion

Using phantoms, structural MRI, EEG, MEG, and mobile electrophysiology, the present studies challenged a foundational assumption underlying non-invasive electrophysiology: that EEG and MEG signals uniquely reflect neural generators embedded in a passive and static volume conductor. Present findings suggest that this assumption is incomplete. Conductive fluid motion alone generated structured, oscillatory, and source-localizable electromagnetic signals in the absence of neural tissue, and interindividual differences in cerebrospinal fluid (CSF) geometry systematically moderated electrophysiological signal amplitude in both EEG and MEG recordings. Crucially, this modulation generalized across participant samples, EEG and MEG recording modalities, sensory paradigms, resting-state activity, and naturalistic locomotion.

The most striking observation was that parietal CSF volume systematically scaled signal amplitude not only in EEG, where conductivity-related effects are expected, but in MEG, which is commonly assumed to be comparatively insensitive to volume-conductor geometry. Larger CSF volumes predicted stronger alpha-band activity, stronger visual gamma responses, and stronger auditory steady-state responses across independent MEG datasets even after controlling for gray matter fraction, age, sex, and dataset identity. The convergence across EEG and MEG argues against a purely modality-specific conductivity explanation. It suggests that CSF geometry contributes more fundamentally to the macroscopic current loops underlying both electric and magnetic field measurements.

Mechanistically, EEG and MEG sensors do not measure neural firing directly but rather electromagnetic fields arising from large-scale current loops(9, 10). Neural transmembrane currents provide the biological drivers of these loops, but current closure necessarily occurs through the surrounding conductive medium. Because CSF forms a continuous, highly conductive compartment surrounding cortical and subcortical structures, variations in CSF geometry may alter the spatial extent and coherence of return-current pathways. In this framework, larger CSF compartments do not merely amplify preexisting neural signals passively but reshape the physical geometry through which macroscopic fields emerge.

Existing finite-element modeling work provides an important reference point for interpreting these findings. In static EEG forward models, CSF is commonly treated as a highly conductive compartment that redistributes return currents and can shunt current away from the scalp, often reducing EEG amplitudes while altering scalp topography(33). The present results differ in both anatomical definition and direction of effect. Here, CSF was quantified as regional parietal CSF volume derived from individual MRI segmentation, rather than as local CSF thickness above a single modeled source, and larger parietal CSF volume was associated with larger, not smaller, EEG and MEG amplitudes. This positive scaling is therefore difficult to explain by a simple static shunting account alone. Instead, it suggests that regional CSF geometry influences the spatial coherence, closure, or mechanical modulation of macroscopic current loops in ways not captured by conventional static forward models. Thus, present findings do not contradict the established importance of CSF in volume conduction, but they indicate that CSF should be considered not as a passive attenuating layer but as a spatially extended and mechanically dynamic conductive compartment.

The present phantom experiments provide an important proof of principle in this regard. Rhythmic displacement of conductive saline generated oscillatory signals with stable topographies and plausible source reconstructions despite the absence of neural tissue. Conventional inverse modeling localized these signals to anatomically plausible cortical regions, demonstrating that realistic source maps can emerge from conductive fluid dynamics alone. These findings do not imply that physiological EEG or MEG signals are non-neural. Rather, they indicate that standard source localization procedures do not uniquely distinguish neural from mechanically induced current loops when both generate coherent macroscopic field structure.

A plausible physical mechanism for the potentials generated in conductive-fluid phantoms is electrokinetic coupling at fluid-surface interfaces(34) (https://en.wikipedia.org/wiki/Double_layer_(surface_science)). When an aqueous electrolyte contacts a charged surface, redistribution of dissolved ions forms an electrical double layer (EDL), comprising relatively immobile surface-associated charge and a mobile counterion-rich diffuse layer. Relative motion of the electrolyte and surface, for example during pressure-driven flow, pumping, sloshing, or tissue deformation, can transport this excess mobile charge, generating a *streaming current* and an associated *streaming potential*. Conversely, the reciprocal electrokinetic phenomenon, *electro-osmosis* (https://en.wikipedia.org/wiki/Electro-osmosis), occurs when an electric field acts on the mobile charge of the EDL and thereby drives fluid motion(34). Thus, electrical activity and fluid dynamics can in principle be coupled bidirectionally through the same charged interface. These mechanisms are well established in electrokinetic systems and hydrated biological tissues, where mechanically induced interstitial-fluid motion can generate streaming potentials and applied electric fields can generate electroosmotic fluid transport(35).

Importantly, such electrokinetic coupling has been experimentally demonstrated in brain tissue, where electrolyte-filled extracellular spaces are bounded by charged cellular surfaces and an extracellular matrix(36, 37). Similar physics may therefore operate in vivo, with CSF and interstitial fluid interacting with cell membranes, extracellular space, vascular basement membranes, and glial interfaces(38-40). Indeed, intracranial-fluid models have explicitly proposed that the high surface-area-to-volume ratio of brain extracellular spaces could support electro-osmotic and other electrokinetic transport and that brain pulsatility and tissue deformation could produce charge motion within the extracellular space(41). In this framework, cardiac pulsation, respiration, vasomotion, cellular volume changes, and glymphatic/perivascular fluid displacement could perturb interfacial charge distributions and generate streaming currents, streaming potentials, and diffusion potentials; conversely, endogenous electrical fields could influence local fluid transport through electro-osmosis.

Such coupling provides a candidate mechanism for the present phantom observations, in which conductive-fluid motion generated structured EEG-like potentials and MEG-detectable magnetic fields in the absence of neural tissue. The phantom experiments do not yet identify the microscopic mechanism or establish the magnitude of such electrokinetic contributions in vivo, but they demonstrate that mechanically driven conductive-fluid dynamics can generate electromagnetic signals measurable with conventional M/EEG instrumentation.

In the brain, such electrokinetic coupling could additionally be driven by activity-dependent cellular volume dynamics(38, 40). Astrocytes actively regulate extracellular ionic and fluid homeostasis, and neuronal activity produces changes in extracellular K^+^, Ca^+^, Mg^+^, and other solutes that are accompanied by water transport, astrocytic swelling, and corresponding changes in extracellular-space volume(38). These processes create motion of ionic extracellular fluid with respect to cell membranes, extracellular matrix, vascular interfaces, and astrocytic processes. Consequently, neural activity could initiate not only conventional transmembrane ionic currents but secondary mechanical and fluid displacements capable, in principle, of generating streaming or diffusion potentials. Such a mechanism would predict that a mechanically generated electromagnetic component could remain tightly coupled to neuronal activity rather than appearing as an independent artifact. Astrocytes are particularly relevant in this context, because they couple neuronal signaling, ionic homeostasis, water transport, extracellular-space geometry, and vascular/perivascular interfaces across spatial scales(7, 38). Recent work further establishes astrocytes as active regulators of neuronal network state rather than passive structural support, with experimental evidence demonstrating activity-dependent astrocyte and extracellular-space volume changes(42-44). Whether these activity-dependent fluid and tissue movements generate electrokinetic currents of sufficient magnitude to contribute measurably to EEG or MEG remains to be established.

The mobile EEG walking study extended these observations into naturalistic behavior. During locomotion, electrophysiological activity tracked individual gait frequency, localized preferentially to sensorimotor and parietal regions adjacent to large CSF compartments, and generated source-reconstructed patterns resembling canonical neural oscillations. Importantly, these rhythmic components scaled with locomotor cadence rather than established endogenous oscillatory frequencies. Although such activity is traditionally treated as motion artifact(45), present findings suggest that movement-related electrophysiological signals instead reflect structured interactions between neural activity, CSF displacement, and mechanically constrained return-current geometry. Supporting this interpretation, the phantom experiment in Fig 1 E-F demonstrated that a single external mechanical perturbation inducing fluid motion in a conductive volume generated sustained oscillatory activity, stable topography, and source localization despite the absence of neural tissue or an internal source generator. Recent work further demonstrates that the brain itself undergoes coherent, behavior-dependent mechanical displacements relative to the skull that contribute to cerebrospinal fluid movement(46), reinforcing the view that electromagnetic activity is embedded within a mechanically dynamic conductive environment rather than a static volume conductor. Moreover, new evidence further indicates that large-scale neural activity is organized as coordinated rotating waves spanning cortical and subcortical systems(47). The present findings suggest that the measured electrophysiological representation of such distributed neural dynamics may also be shaped by the mechanical and conductive properties of the surrounding CSF compartment. This issue is increasingly relevant as neuroscience moves toward mobile and wearable electrophysiology in real-world settings.

In summary, present findings do not deny the central role of neural activity in generating EEG and MEG signals. Neural currents remain necessary drivers of macroscopic electromagnetic fields. Rather, the results indicate that the surrounding conductive medium contributes actively to how those fields emerge, propagate, and are measured. Across phantom experiments, EEG, MEG, and mobile recordings, CSF geometry systematically shaped measured signal amplitude and source structure, suggesting that the physical configuration and dynamics of the conductive medium are integral determinants of the electromagnetic fields ultimately recorded at the scalp. The present work supports a reframing of non-invasive bioelectromagnetism as a measurement of neural activity embedded in a mechanically active conductive system rather than a static mapping from neural generators to sensors.

## Acknowledgments

The authors thank Nathan Weisz and Gianpaolo Demarchi for generously providing access to the MEG facility and for their support with the MEG phantom experiments at the Centre for Cognitive Neuroscience, University of Salzburg. The authors also acknowledge the members of the Heller/Miller lab at the University of Illinois Urbana-Champaign for their contributions to the human MRI/EEG dataset(14) used for new analyses reported here. T.P. used ChatGPT (OpenAI) during the preparation of early manuscript drafts solely for grammar editing and language refinement. No AI tool was used for study design, data analysis, interpretation of results, or generation of scientific content. G.A.M. did not use any AI-assisted technologies. All scientific content, analyses, interpretations, and conclusions were conceived, verified, and approved by the authors.

## Data and code availability

Processed data underlying all analyses, the phantom datasets, and the analysis code will be made publicly available through the Open Science Framework (OSF) upon publication. The human MRI+EEG dataset cannot be shared in raw form because participant consent and institutional ethics approval do not permit unrestricted public release of identifiable electrophysiological recordings. Processed MRI+EEG data necessary to reproduce the reported analyses will be provided through the OSF repository. The remaining datasets analyzed in this study (including publicly available MEG datasets) are available from their original repositories as cited in the manuscript.

